# A real-time colorimetric reverse transcription loop-mediated isothermal amplification (RT-LAMP) assay for the rapid detection of highly pathogenic H5 clade 2.3.4.4b avian influenza viruses

**DOI:** 10.1101/2023.05.14.540682

**Authors:** Fabien Filaire, Aurélie Sécula, Laetitia Lebre, Guillaume Croville, Jean-Luc Guerin

## Abstract

Highly pathogenic avian influenza viruses (HPAIV) are a major threat to the global poultry industry and public health due to their zoonotic potential. Since 2016, Europe and France have faced major epizootics caused by clade 2.3.4.4b H5 HPAIV. To reduce sample-to-result times, point-of-care testing is urgently needed to help prevent further outbreaks and the propagation of the virus. This study presents the design of a novel real-time colorimetric reverse transcription loop-mediated isothermal amplification (RT-LAMP) assay for the detection of clade 2.3.4.4b H5 HPAIV. A clinical validation of this RT-LAMP assay was performed on 198 pools of clinical swabs sampled in 52 poultry flocks during the H5 HPAI 2020-2022 epizootics in France. This RT-LAMP assay allowed the specific detection of HPAIV H5Nx clade 2.3.4.4b within 30 minutes with a sensitivity of 86.11%. This rapid, easy-to-perform, inexpensive, molecular detection assay could be included in the HPAIV surveillance toolbox.

## Introduction

Avian influenza viruses (AIV) are enveloped negative-sense segmented single-stranded RNA viruses belonging to the *Orthomyxoviridae* family. AIV are commonly classified into subtypes based on the combination of their surface glycoproteins, hemagglutinin (HA) and neuraminidase (NA) (Wahlgren, 2011). Importantly, AIV also can be differentiated by their pathogenicity: low pathogenic avian influenza viruses (LPAIV) and highly pathogenic avian influenza viruses (HPAIV). LPAIV are the most predominant worldwide and cause no to mild symptoms in infected individuals (Germeraad *et al*., 2019; Jourdain *et al*., 2010). Most of the time, LPAIV are undetected in flocks. However, due to genetic changes, H5 and H7 HA sometimes can shift from low pathogenic to highly pathogenic forms (Abdelwhab *et al*., 2013; Dupré *et al*., 2021; Wahlgren, 2011). Unlike LPAIV, HPAIV induce severe disease associated with strong typical clinical signs and high mortality rates (Lean *et al*., 2022).

HPAIV display a tremendous evolutionary potential, driven by mutations, indels and reassortments (D. Lee *et al*., 2017; D. H. Lee *et al*., 2021; L. Shi *et al*., 2019). This property has led to the emergence of the A/goose/Guangdong/1996(Gs/GD) lineage. All subsequent viruses derived from this lineage have been classified into clades and subclades. Over the years, clade 2.3.4 has become dominant, and subclade 2.3.4.4b has been widely persistent in Eurasia since 2016 (W. Shi & Gao, 2021). Viruses belonging to clade 2.3.4.4b have led to major epizootics, mostly due to H5N8 subtypes from 2016/2017 to 2020/2021, and then an H5N1 subtype in 2021/2022. HPAIV epizootics have caused the death of millions of wild and domestic birds, threatened public health due to zoonotic risks, and generated major economic losses to the poultry industry (Adlhoch *et al*., 2022). In addition, HPAIV epizootics seem to be spreading more widely over time, with new territories infected worldwide.

HPAIV infections are characterized by a massive viral shedding in the early stages of the infection, especially in ducks (Gaide *et al*., 2021; Vergne *et al*., 2021), and a rapid spread of the disease. These findings raise numerous challenges for the control and the surveillance of the infection, especially during epizootics. Despite the reinforcement of biosecurity measures since 2016 (Delpont *et al*., 2021) and the application of new control measures by public authorities each year, the virus has acquired the ability to spread rapidly and uncontrollably (Guinat *et al*., 2020; Lewis *et al*., 2021; Vergne *et al*., 2021). New strategies for the early detection of HPAIV, therefore are needed to better control viral spread.

Currently, European and French official guidelines for HPAIV detection and surveillance require an rRT-PCR analysis on tracheal swabs (Nielsen *et al*., 2021). Positive results for HPAIV systematically lead to the culling of entire poultry flocks. However, although rRT-PCR is considered as the gold standard for HPAIV detection based on its analytical sensitivity, this method is still rather expensive, time-consuming (∼80 min), and requires sophisticated equipment that is difficult to transport and must be operated by trained staff. This ultimately reduces its in-field diagnosis potential and can delay the rapid response needed during an epizootic.

Easy-to-perform, fast, low-cost, low-tech, sensitive, and specific methods are needed to improve the rapidity of HPAIV detection and develop new strategies for HPAIV surveillance in the field. The loop-mediated isothermal amplification (LAMP) assay is a molecular biology technology that has been developed since 2000 (Notomi *et al*., 2000). The LAMP assay is an end-point nucleic acid amplification method based on a DNA polymerase. The technology requires a set of 2 or 3 pairs of primers targeting 6 to 8 binding sites, making it highly specific. The fast and isothermal amplification (15-40 min) can be performed by standard transportable equipment such as a heat block or water bath. This, combined with user-friendly read-outs such as fluorescence, turbidity and even colorimetric changes, enables its utilization in a field point-of-care strategy. Overall, LAMP assays are inexpensive (∼1.5 euros/reaction, for indication of magnitude only), rapid, easy-to-perform, and robust against the well-known PCR reaction inhibitors. For all of these reasons, LAMP assays have been largely developed for the detection of viruses (Golabi *et al*., 2021; Padzil *et al*., 2022; Vanhomwegen *et al*., 2021). More importantly, the low-technology required and the possible lyophilization of reagents allow its utilization in remote locations where resources are scarce or non-existent (Howson *et al*., 2017; Kumar *et al*., 2021; Vanhomwegen *et al*., 2021).

Our research focused on developing a real-time colorimetric reverse transcription LAMP (RT-LAMP) assay for the detection of H5 HPAIV clade 2.3.4.4b. We designed a primer set for the detection of H5 HPAIV clade 2.3.4.4b, and assessed its sensitivity and specificity on eight different AIV subtypes and pathotypes, including viruses from diverse clades. Finally, we assessed the performance of this real-time colorimetric RT-LAMP assay on tracheal and cloacal swabs sampled in France during the 2020/2021 and 2021/2022 H5 HPAIV epizootics.

## Materials and Methods

### Primer design

A set of 626 HA sequences from H5 HPAIV clade 2.3.4.4b HA available on GISAID up to 7 February 2022, *via* https://gisaid.org or from our laboratory, were aligned using Geneious Prime 2021.2.2 (https://www.geneious.com). From this alignment, a consensus sequence with a 95% base identity threshold was extracted to target a 200 to 350 base-long region with low base diversity. Finally, a 308 base region was selected and passed through the PrimerExplorer V5 online tool for LAMP primer design. A set of primers was selected (Table 1) and their specificity was checked through a BLAST alignment with eight selected AIV subtypes (Table 2).

**Table 1:**
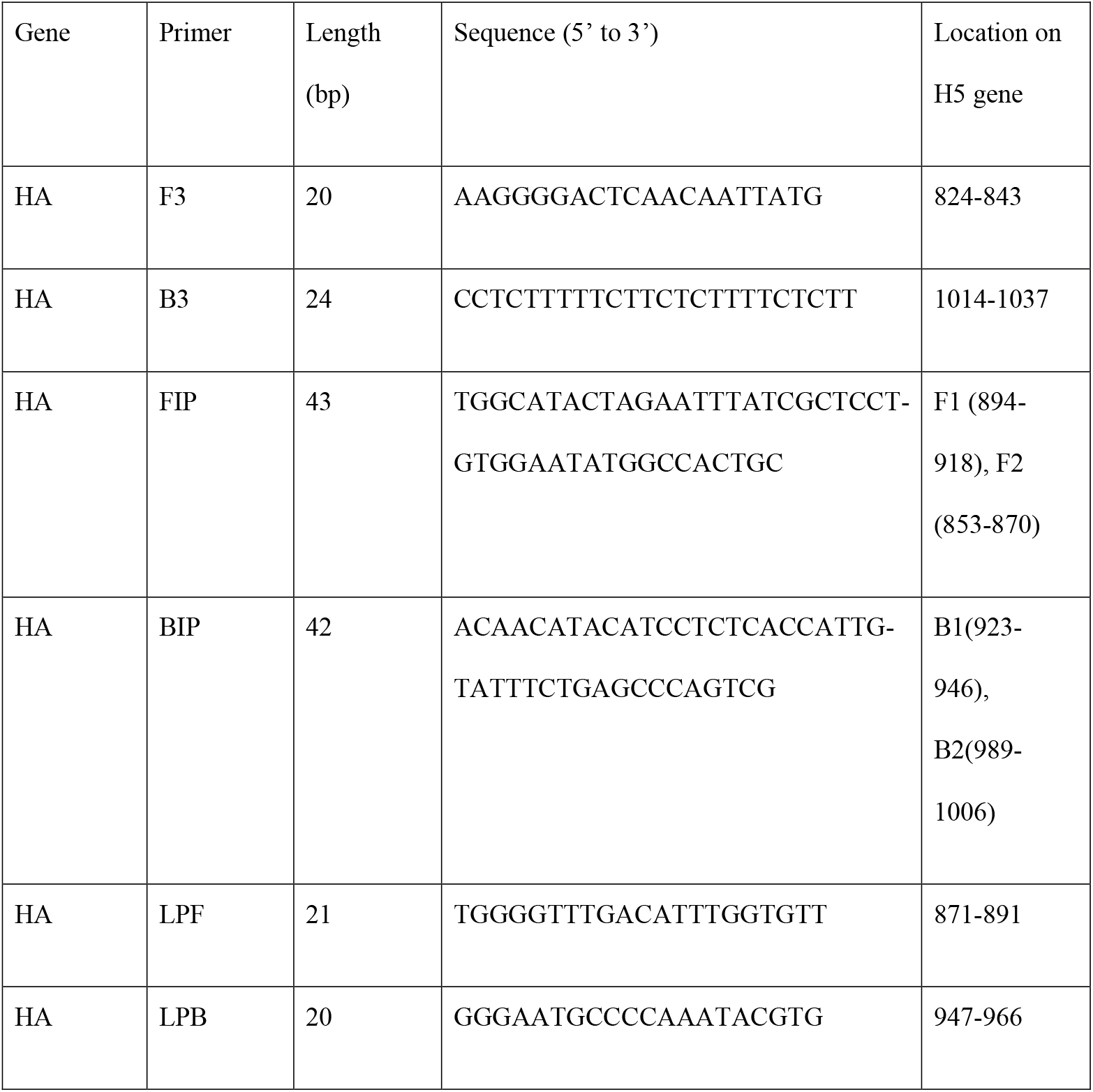
List and sequences of the designed RT-LAMP primers. The nucleotide position corresponds to the HA sequence of the HPAIV H5N8 2020/2021 (accession number MZ166300).

**Table 2:** Viruses selected to test the designed primer set

| Subtype | Pathogenic form | Clade 2.3.4.4b | Sampling location | Sampling time | Host | Amplification | Virus reference | Accession number |
| --- | --- | --- | --- | --- | --- | --- | --- | --- |
| H5N8 | HP | Yes | France | 2016 | Duck | Embryonated eggs | A/mallard duck/France/171201g/2017 (H5N8) | MN875092 |
| H5N8 | HP | Yes | France | 2020 | Duck | Embryonated eggs | A/Mule_duck/France/20353/2020(H5N8) | MZ166300 |
| H5N1 | HP | Yes | France | 2021 | Duck | Embryonated eggs | A/Mule_duck/France/21348/2021 (H5N1) | OP828913 |
| H5N9 | HP | No | France | 2015 | Guinea Fowl | Embryonated eggs | A/Guinea Fowl/France/129/2015(H5N9) | MN400996 |
| H9N2 | LP | No | Tunisia | 2021 | Chicken | None | A/Gallus gallus/Tunisia/20057/2020(H9N2) | OP828915 |
| H6N1 | LP | No | France | 2021 | Duck | Embryonated eggs | A/Pekin_duck/France/21114/2021(H6N1) | OP828914 |
| H7N1 | LP | No | Italy | 1999 | Turkey | Cells (MDCK) | A/Turkey/Italy/977/1999/H7N1 | GU052999 |
| H5N3 | LP | No | Italy | 2004 | Duck | Embryonated eggs | A/duck/Italy/775/2004(H5N3) | CY024746 |

### Inclusivity and exclusivity

The analytical inclusivity and exclusivity of the designed primers were tested *in silico*, by rRT-PCR and by our RT-LAMP assay, on eight different avian influenza subtype viruses available in our laboratory. The selection included three H5 HPAIV clade 2.3.4.4b, one H5 HPAIV from the European lineage (not part of clade 2.3.4.4b) (Briand *et al*., 2017) and four LPAIV (Table 2).

### RNA extraction

Viral RNA was extracted using the magnetic bead-based ID Gene Mag Fast Extraction Kit (IDvet, Grabels, France) combined with the IDEAL 32 extraction robot (IDvet), following the manufacturer’s instructions. The extracted RNA was stored at -20 °C before use.

### Colorimetric Real-time RT-LAMP

RT-LAMP reactions were done using the WarmStart Colorimetric LAMP 2X Master Mix kit (M1800, NEB, Hitchin, UK) following the manufacturer’s instructions. Briefly, a 25 μL reaction mix was prepared by mixing 12.5 μL of the WarmStart Colorimetric LAMP 2X Master Mix, 5 μL of the sample viral RNA, 5 μL nuclease free water and 2.5 μL of a 10x primer solution mix prepared beforehand. The 10x primer solution mix was prepared with 16 μM of each FIP/BIP, 2 μM of each B3/F3, and 4 μM of each LF/LB. The reaction mix was incubated at 65 °C for 30 min and the colour switch, from purple to yellow in case of positive reaction, was assessed with the naked eye.

### Real-time RT-qPCR

Two quantitative RT-PCR were used in this study. A real-time quantitative RT-PCR (rRT-PCR) with the officially-approved IDvet M gene and H5/H7 one-step rRT-PCR kit (Idvet, https://www.id-vet.com) was used for clinical validation as the ‘gold standard’ molecular-based HPAIV detection method. All reactions were realized following the manufacturer’s instructions. A second RT-qPCR was used for the limit of detection assay. Viral RNA absolute quantification was performed using the iTaq Universal SYBR green one-step kit (#1725150, Bio-rad). The H5 gene was targeted with HA H5N8_HP_ primers (5’-GACCTCTGTTACCCAGGGAGCCT-3’, 5’-GGACAAGCTGCGCTTACCCCT-3’) (Bessière *et al*., 2021). The absolute quantification was performed using a standard curve based on 10-fold serial dilution of a plasmid containing the H5 HA gene. RT-qPCR reaction and results analysis were performed on a LightCycler 96 instrument (Roche).

### Limit of detection assay

To investigate the detection limit of this RT-LAMP assay, a viral RNA absolute quantification was performed. Two serial dilutions of extracted viral RNA from embryonated eggs HPAIV H5N8 2020/2021 and HPAIV H5N1 2021/2022 amplification were realized. Each dilution was systematically analyzed by our RT-LAMP assay and the iTaq RT-qPCR.

### Clinical validation

The designed real-time colorimetric RT-LAMP was tested for clinical use on swabs in an epizootic context. A total of 198 swabs pools, corresponding to 52 poultry flocks (32 duck, 19 chicken, 1 quail flocks), was included in this validation (Supplementary Table 1). The pools consisted of a mix of five tracheal or cloacal swabs sampled in infected or suspected farms during the HPAIV H5N8 2020/2021 and HPAIV H5N1 2021/2022 epizootics in France. All samples were tested in accordance with the European guidelines for HPAIV diagnosis. First, the total RNA was extracted using the magnetic bead-based ID Gene Mag Fast Extraction Kit IDvet (https://www.id-vet.com) combined with the IDEAL 32 extraction robot (IDvet), following the manufacturer’s instructions. Then, HPAIV H5Nx viral RNA detection was simultaneously performed by rRT-PCR with the IDgene H5/H7 one-step rRT-PCR kit and the real-time colorimetric RT-LAMP assay. To validate the feasibility of the protocol under field conditions, this clinical validation was performed by a non-trained member of staff.

## Results

The designed RT-LAMP primers were analyzed *in silico* to confirm their complementarity to binding regions. Multiple alignments with publicly available HA sequences from different AIV subtypes, including HPAIV from different clades and LPAIV, were selected. The alignment revealed the high binding complementary to the targeted regions of the clade 2.3.4.4b H5 HPAIVs only. The overall binding complementary reaches a 97.64%, 100%, and 99.4% ratio for the HPAIV H5N8 2016/2017, 2020/2021 and the HPAIV H5N1 2021/2022, respectively. However, for the non-clade 2.3.4.4b viruses, the overall complementary rates do not exceed 84.12% (Supplementary Table 2). This low binding affinity is theoretically not sufficient to allow amplification (26,27) (Figure 1). To investigate the analytical inclusivity and exclusivity of the different AIV subtypes, eight AIV from distinct subtypes and pathotypes, available in our laboratory, were used (Table 2, Figures 1 and 2). All samples were analyzed using the RT-LAMP assay in parallel with the gold standard rRT-PCR targeting both the M, H5 and H7 genes as controls for the detection of viral RNA. The results showed a 100% (3/3) analytical reactivity for the detection of clade 2.3.4.4b H5 HPAIV. Additionally, the exclusivity test showed a 100% primer specificity as none of the non-H5 clade 2.3.4.4b viruses tested positive (Figure 2). These results, associated with the *in silico* analysis, tend to confirm the specificity of the RT-LAMP assay for HPAIV H5Ny from clade 2.3.4.4b.

**Figure 1.**
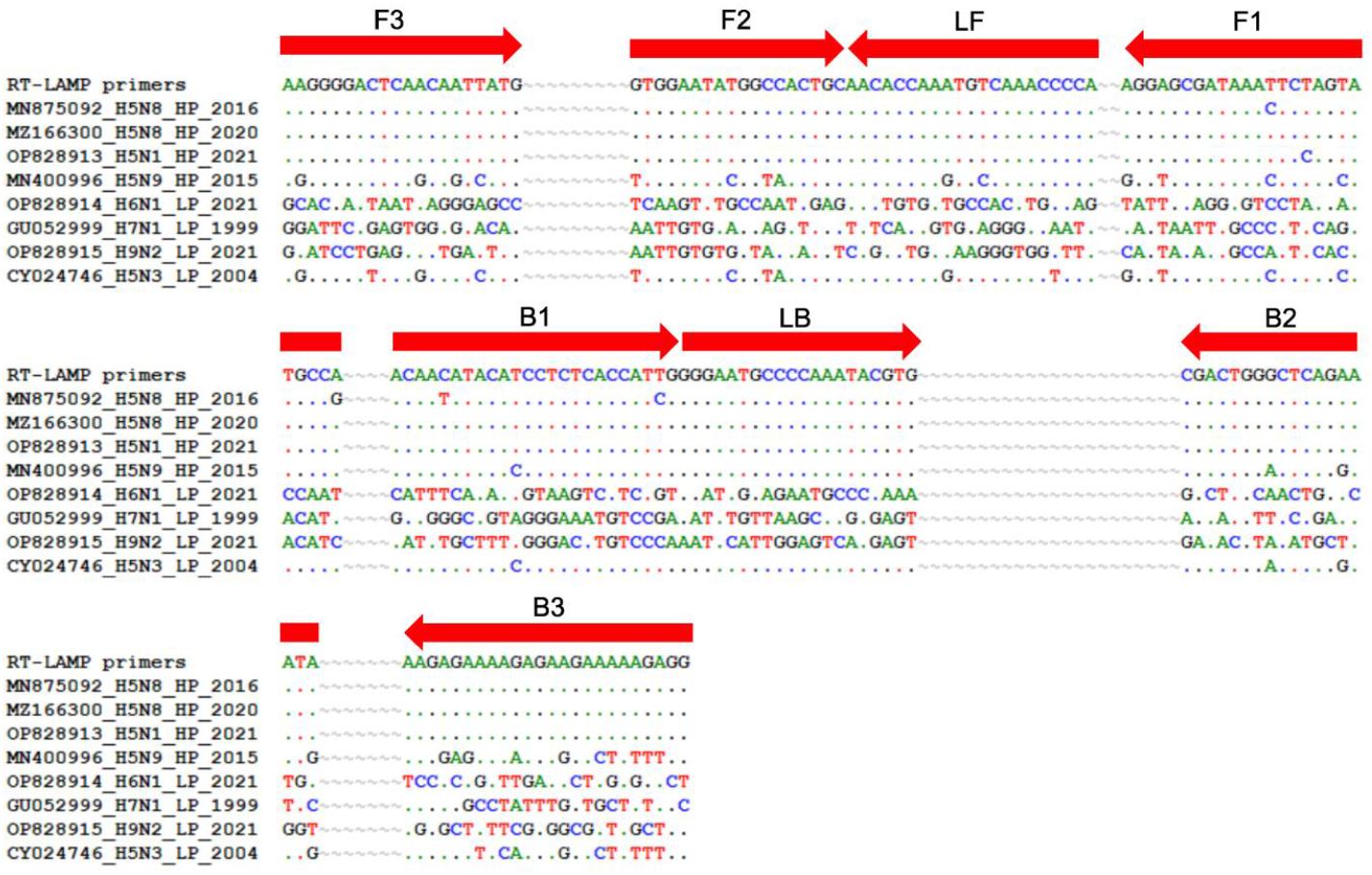
LAMP primers alignment to the sequence of the 8 AIV viruses included in the study.

**Figure 2.**
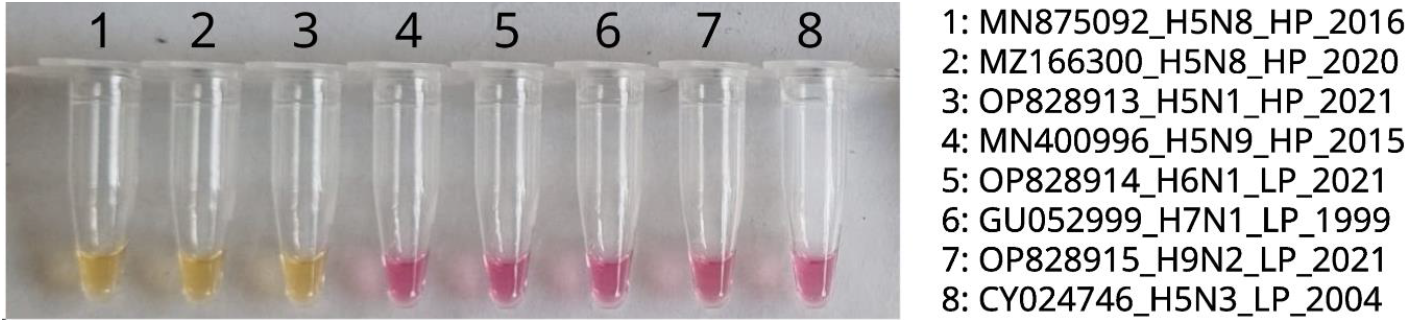
Real-time colorimetric LAMP results on different AIV viruses. Positive amplification induces a colorimetric change from purple to yellow. 1, 2016/2017 H5N8 HP clade 2.3.4.4b virus. 2, 2020/2021 H5N8 HP clade 2.3.4.4b virus. 3, 2021/2022 H5N1 clade 2.3.4.4b virus. 4, H5N9 HP none clade 2.3.4.4b virus. 5, H6N1 LP virus. 6, H7N1 LP virus. 7, H9N2 LP virus. 8, H5N3 LP virus.

Furthermore, the RT-LAMP assay detection limit was investigated by absolute quantification of the viral RNA by the iTaq RT-PCR. Two separate 2-fold serial dilutions of HPAIV H5N8 2020/2021 and H5N1 2021/2022 viral RNA samples were analyzed by RT-LAMP assay, rRT-PCR and RT-qPCR (Supplementary Table 3). The results were globally similar for both viruses. Indeed, the lowest RNA concentrations testing positive with the RT-LAMP assay were 9.66 and 18.44 copy /μL for the H5N8 HP 2020/2021 and the H5N1 HP 2021/2022, respectively (Supplementary Table 3). These findings are in agreement with the expected detection limit of a RT-LAMP assay, usually ranging from 100 to 1000 copies/reaction (Zhang *et al*., 2020). Moreover, the rRT-PCR results indicate a shared limit of detection with a cycle threshold (C_t_) of around 30 for both viruses (Supplementary Table 3).

Finally, a clinical validation of this RT-LAMP assay was performed on clinical swabs sampled during the H5N8 HP 2020/2021 and H5N1 HP 2021/2022 epizootics in France. Following the European guidelines for HPAIV surveillance and detection, a total of 198 pools of five tracheal or cloacal swabs each, corresponding to 52 poultry flocks (32 duck, 19 chicken, 1 quail flocks) (Supplementary Table 4), were analyzed simultaneously by rRT-PCR, considered the gold standard method, and by the RT-LAMP assay developed.

Firstly, the swabs were grouped, regardless of the flocks, based on their C_t_ values obtained by rRT-PCR to investigate the sensitivity and specificity of the RT-LAMP assay depending on the viral RNA loads. Therefore, samples were divided for into four categories: C_t_ <25, 25< C_t_ <30, C_t_ >30 and 0<C_t_ (Supplementary Table 4). The results of the rRT-PCR and RT-LAMP showed a very good agreement for samples with C_t_s below 30. Indeed, the RT-LAMP assay showed a 100% and 86.27% sensitivity for samples with C_t_<25 and C_t_s between 25 and 30, respectively. For C_t_ values above 30, the sensitivity decreased drastically to 18.18%. When all data were considered altogether without differentiation based on C_t_s, the overall sensitivity and specificity reached 86.11% and 100%, respectively (Supplementary Table 4).

Secondly, the data were analyzed grouped by flocks. Our data regrouped a total of 52 flocks with between 1 and 9 swab pools. According to the HPAIV detection guidelines (Nielsen *et al*., 2021), only one pool must be detected as positive to consider the whole flock positive. Based on this guidance, only five flocks showed result discrepancies between RT-LAMP and rRT-PCR assays (RT-LAMP assay negative while rRT-PCR positive) (Supplementary Table 1). In this context, data analysis showed a 90% sensitivity and 100% specificity for HPAIV detection in flocks. Interestingly, the five flocks with divergent results correspond to samples with RT-PCR C_t_>30.

## Discussion

To improve the molecular detection of HPAIV circulating worldwide, this study aimed to develop a real-time colorimetric RT-LAMP assay. Taken together, our findings suggest that this new real-time colorimetric RT-LAMP assay may offer, in specific contexts and purposes, an alternative to the gold standard rRT-PCR for the detection of HPAIV from clade 2.3.4.4b provided the primers are regularly updated. The three couples of designed primers have shown to be highly specific to the clade 2.3.4.4b H5 HPAIV in both *in silico* and *in vitro*. However due to the multiplicity of primers binding to a total of 8 binding sites, a high primer-to-binding site complementarity is required. Even though a relative difference of detection limit and sensitivity can be noted between distinct viruses (Table 2, Supplementary Table 3 and 4), this assay has proven to be very sensitive, with a detection threshold determined below 20 copy/ μL. Previous knowledge of the circulating strains (i.e., obtained by sequencing in a context of diagnosis or surveillance) is highly recommended to avoid false negatives due to a lack of specificity. Following the HPAIV detection guidelines, RT-LAMP detection of the M gene could be performed simultaneously (Golabi *et al*., 2021).

A clinical validation performed on 198 pools of clinical swabs sampled in 52 poultry flocks have shown an overall sensitivity of 86.11% with up to 100% for samples with C_t_ below 25. Moreover, even though both detection limit investigations done by RT-qPCR and the clinical assay showed suboptimal results for samples with C_t_>30, this does not seem to be a major limitation in the context of an HPAIV epizootic. Indeed, most HPAIV infections induce high viral shedding, even in the earliest stages of the infection, leading to high loads of viral RNA (Criado *et al*., 2021; Filaire *et al*., 2022; Germeraad *et al*., 2019) especially in the respiratory tract (Gaide *et al*., 2021). Therefore, low viral RNA loads, associated with C_t_>30, which corresponds to approximately less than 20 copies / μl (Table 2 and Supplementary Table 4), are infrequent in HPAIV-infected birds.

This 30 min colorimetric RT-LAMP reaction could be performed in the field for point-of-care application. Additionally, this strategy could be included in a workflow comprising fast lysis and extraction methods and environmental sampling methods for the rapid detection of new outbreaks directly on-farm, especially in a context of clinical suspicions, where viral RNA loads are the highest. Further validation by proficiency tests in reference laboratories would be required before implementation in official surveillance but in the principle, this rapid, easy-to-perform (even for non-trained staff), inexpensive, low-tech molecular detection assay could be included in the HPAIV surveillance toolbox and improve the response capacity during epizootics.

## Acknowledgement

This study was performed in the framework of the Chaire de Biosécurité & Santé Aviaires, hosted by the National Veterinary College of Toulouse (ENVT) and funded by the Direction Generale de l’Alimentation, Ministère de l’Agriculture et de la Souveraineté Alimentaire, France. Fabien Filaire’s PhD position is funded by Theseo France and Lanxess biosecurity from the Lanxess group. The funders had no role in study design, data collection and interpretation, or the decision to submit the work for publication.We would like to thank the farm holders for their collaboration with sampling of their flocks and show our gratitude to Sebastien Mathieu Soubies for comments and constructive criticism on an earlier version of the manuscript. Finally, all authors acknowledge GISAID contributors.

## Supplementary Material

**Supplementary Table 1:**
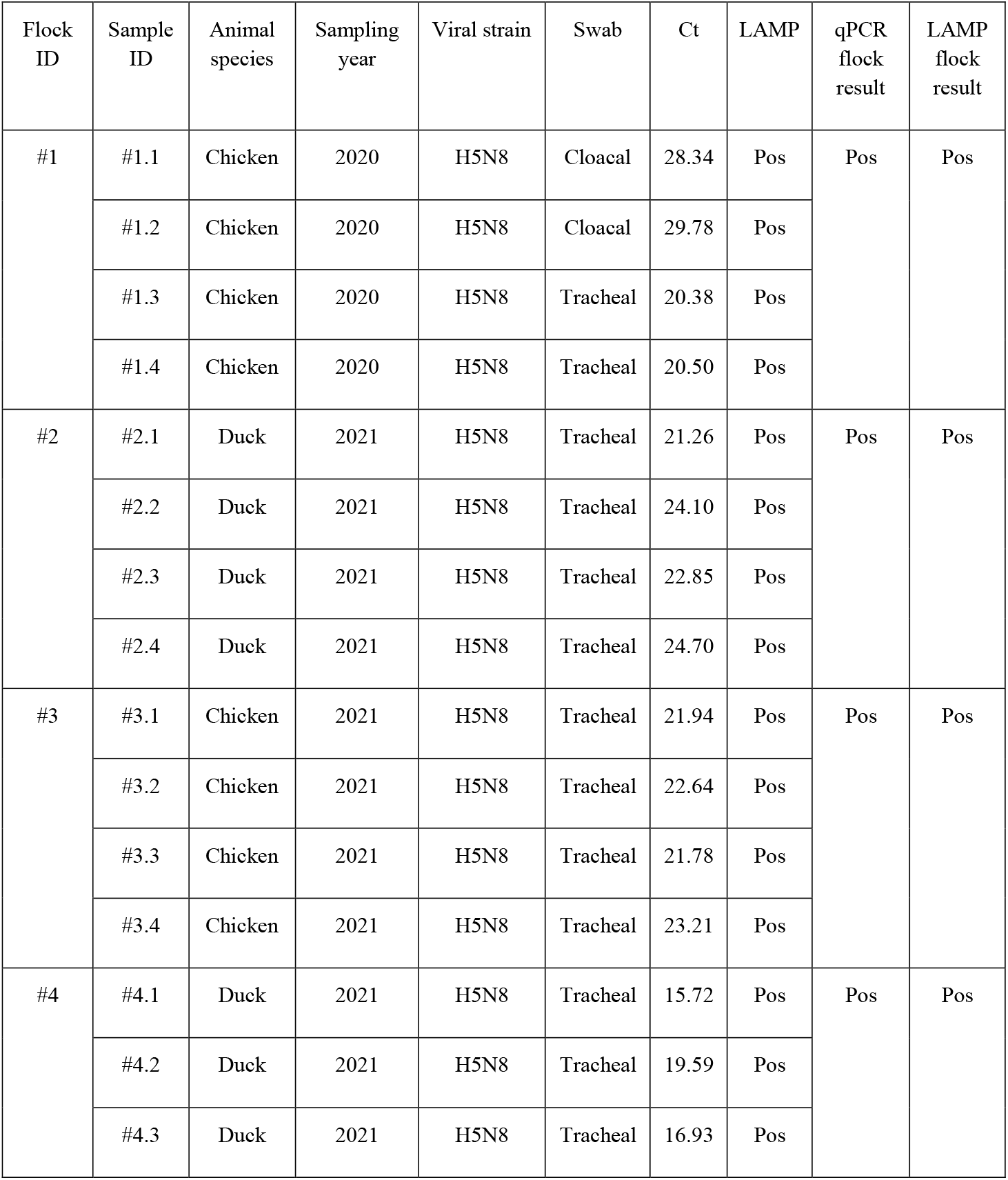
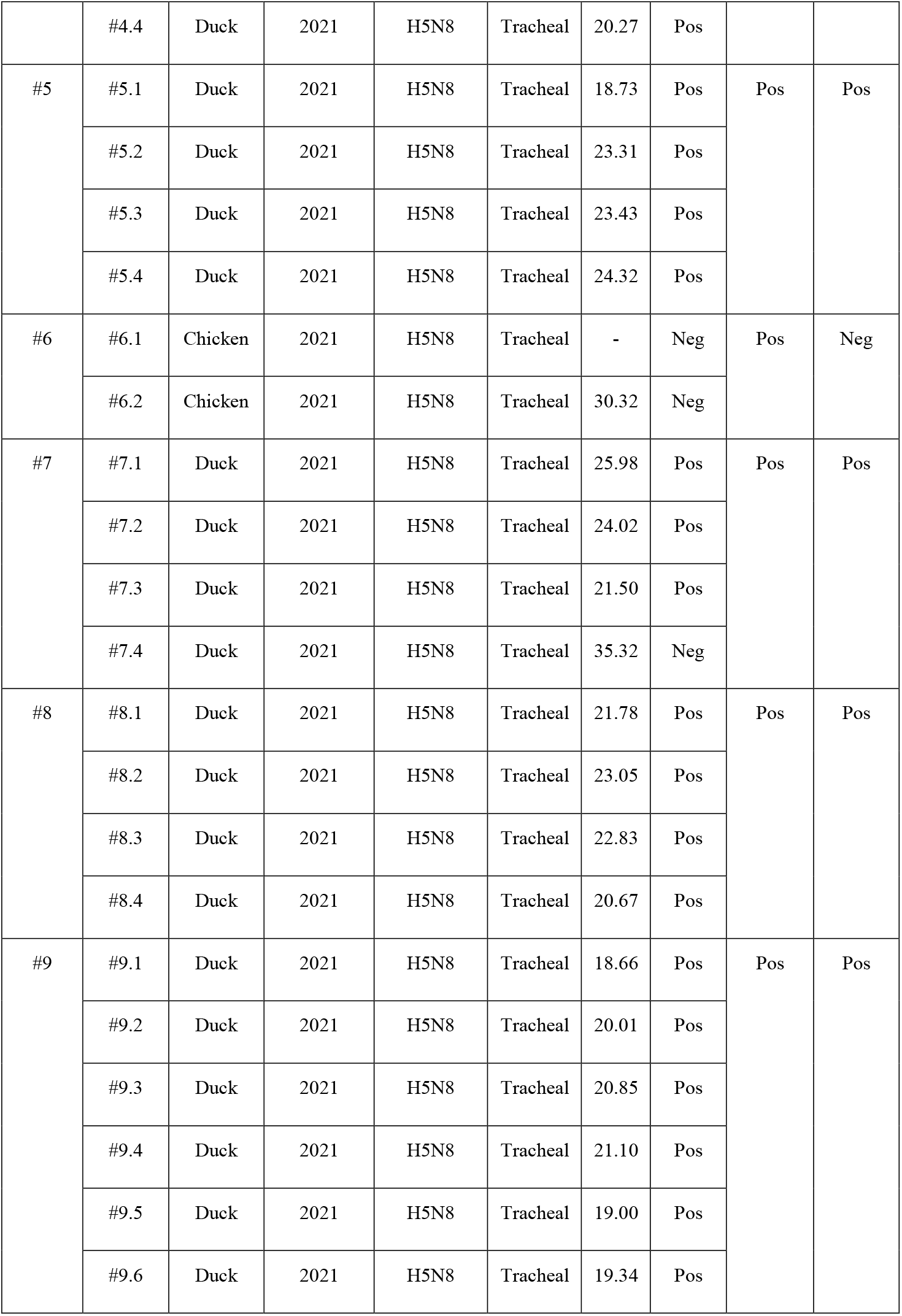
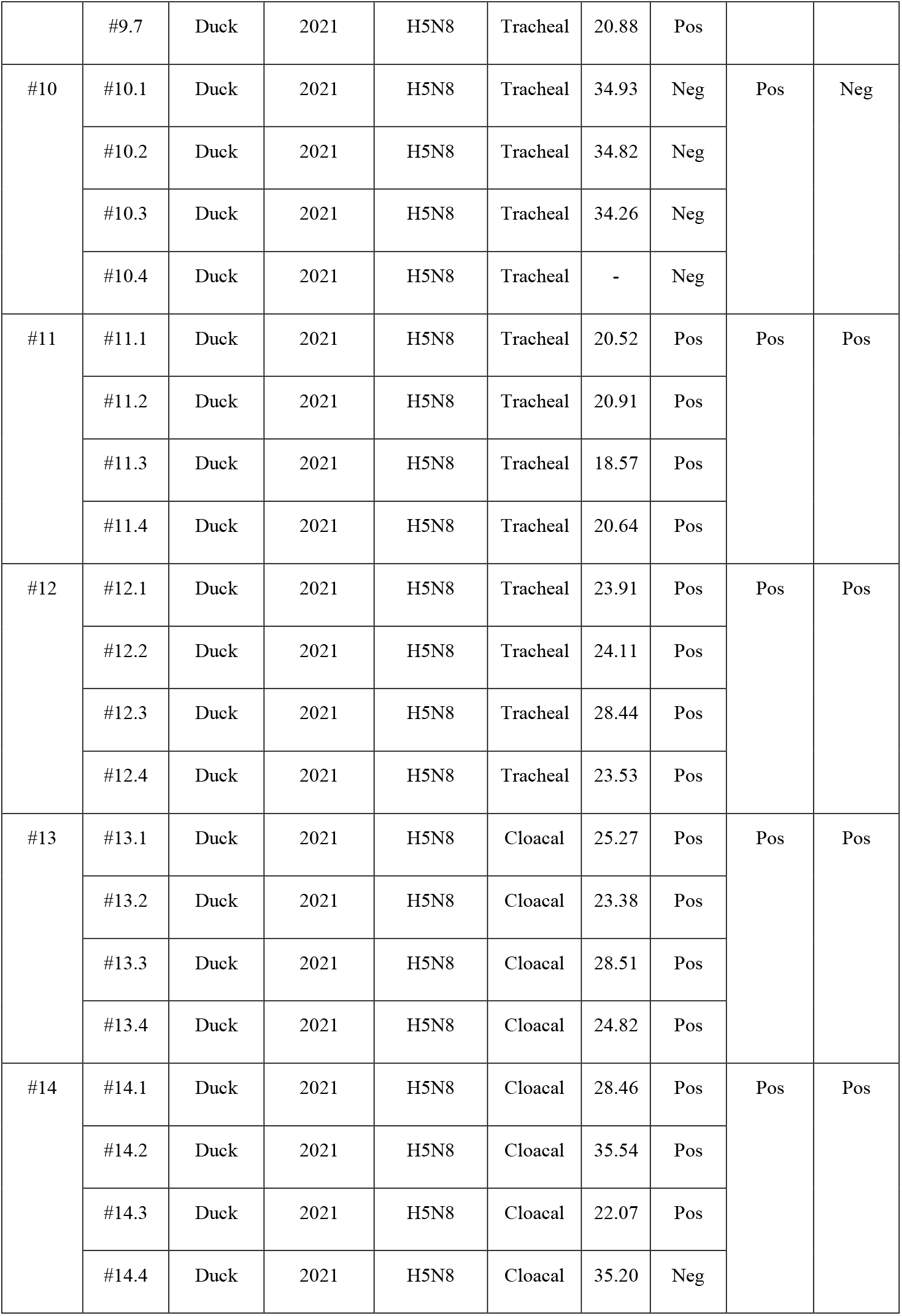
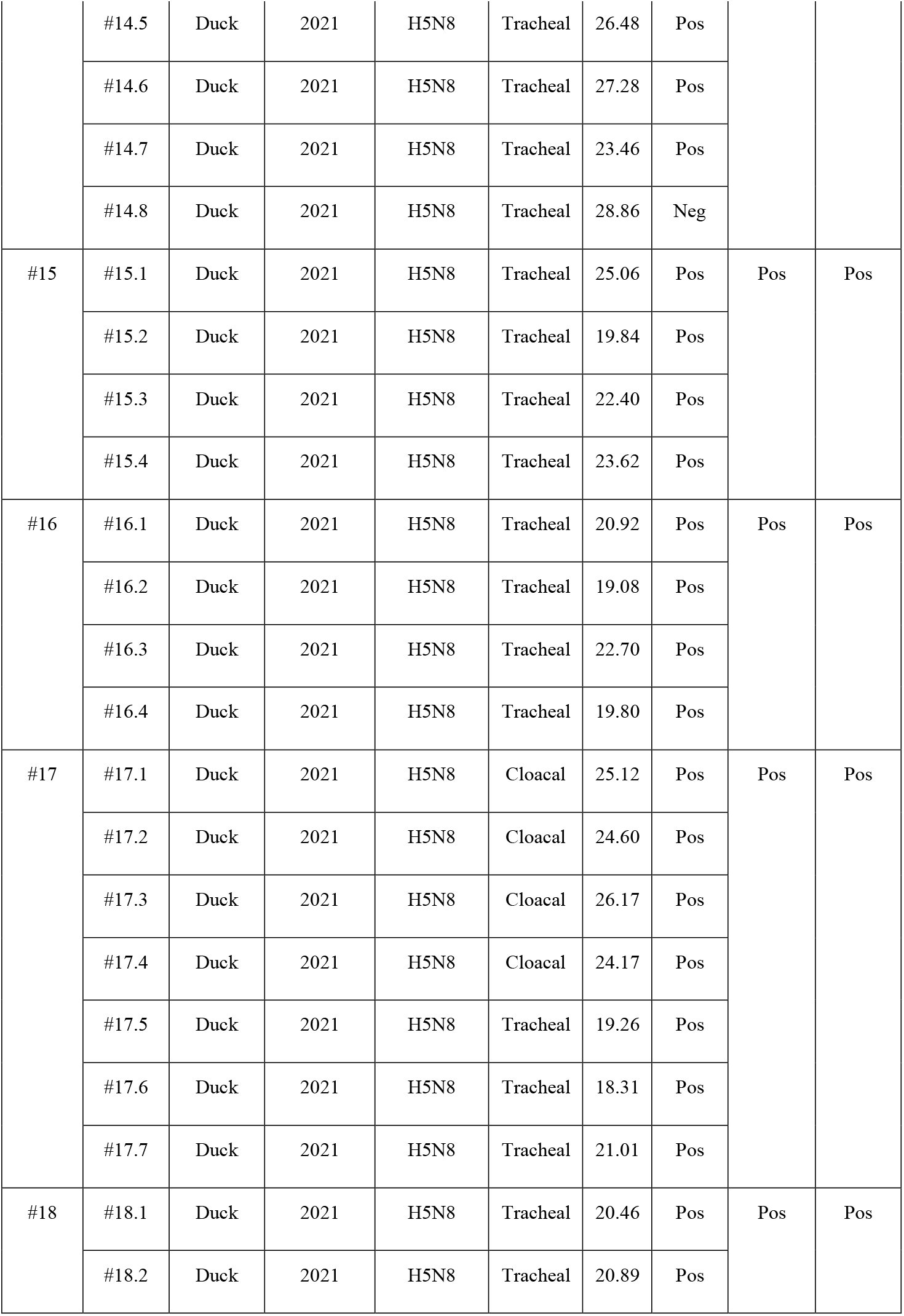
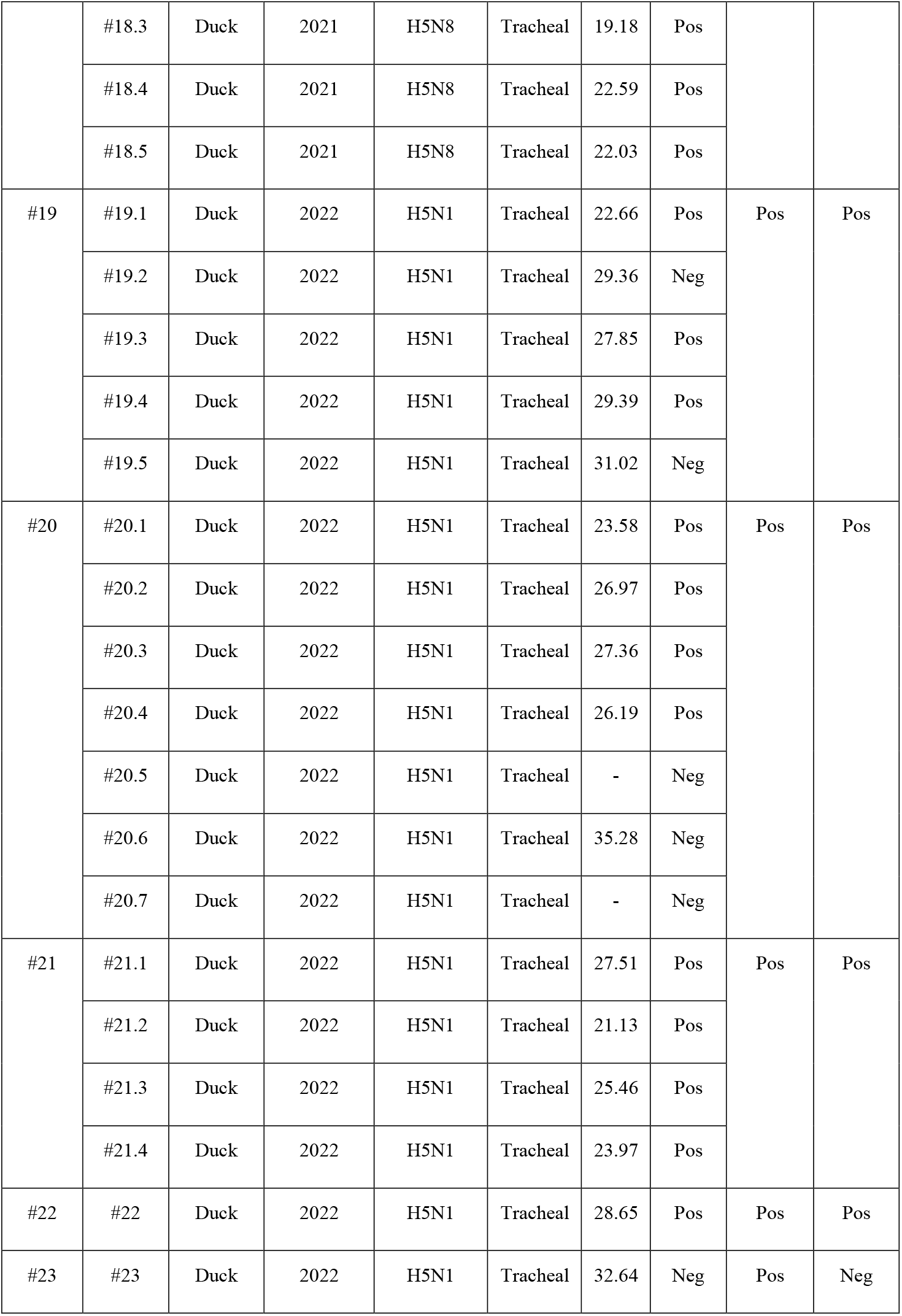
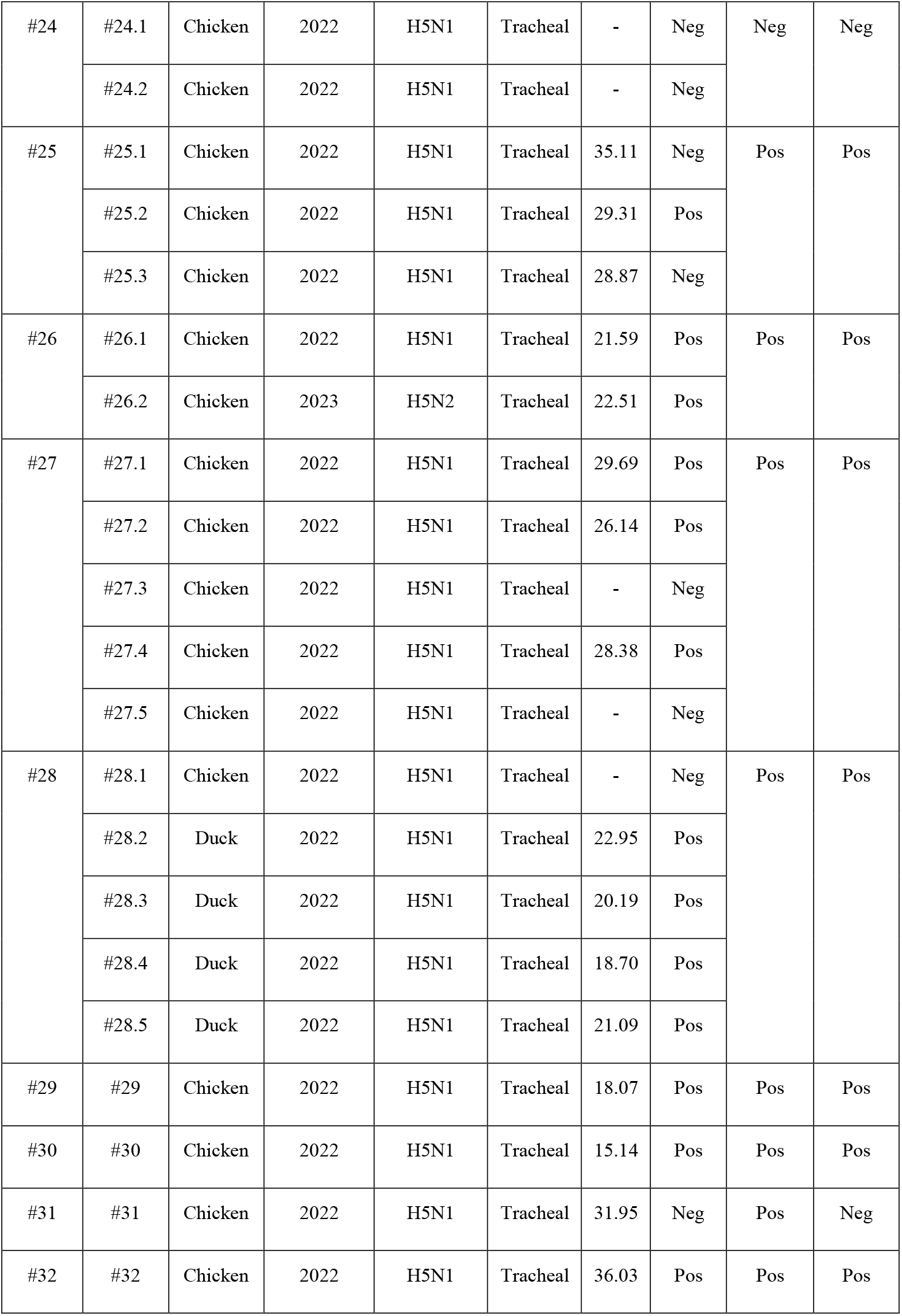
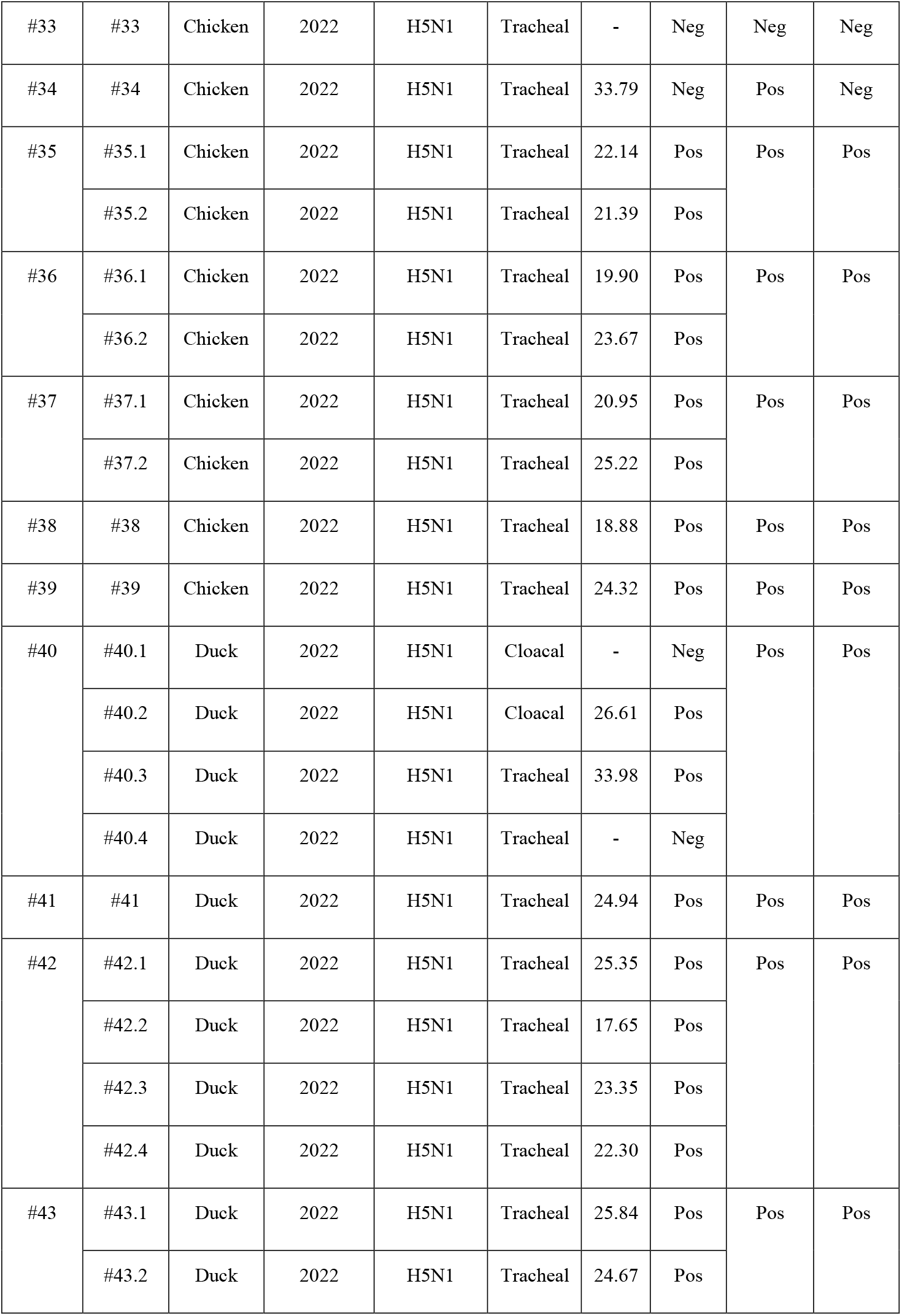
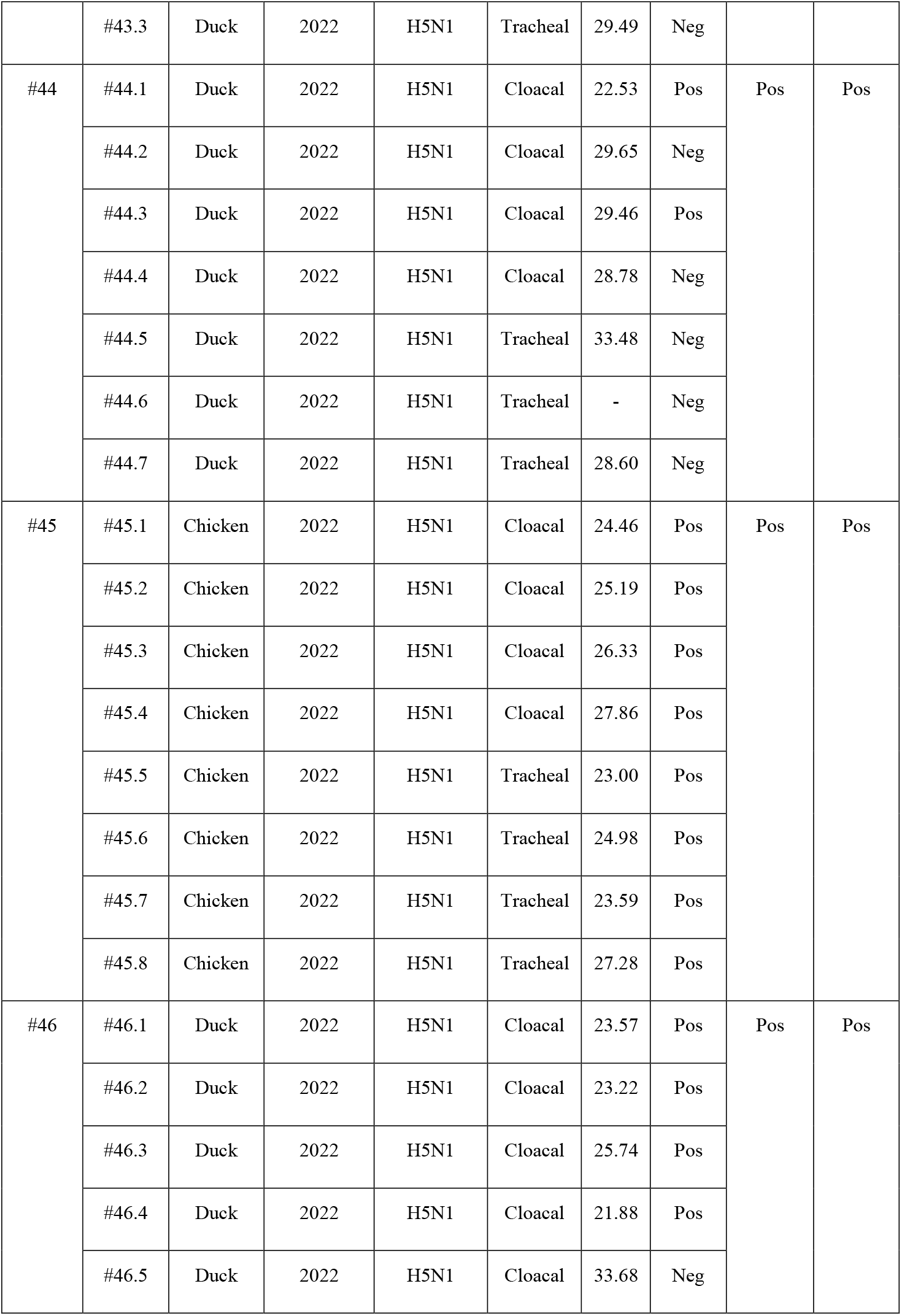
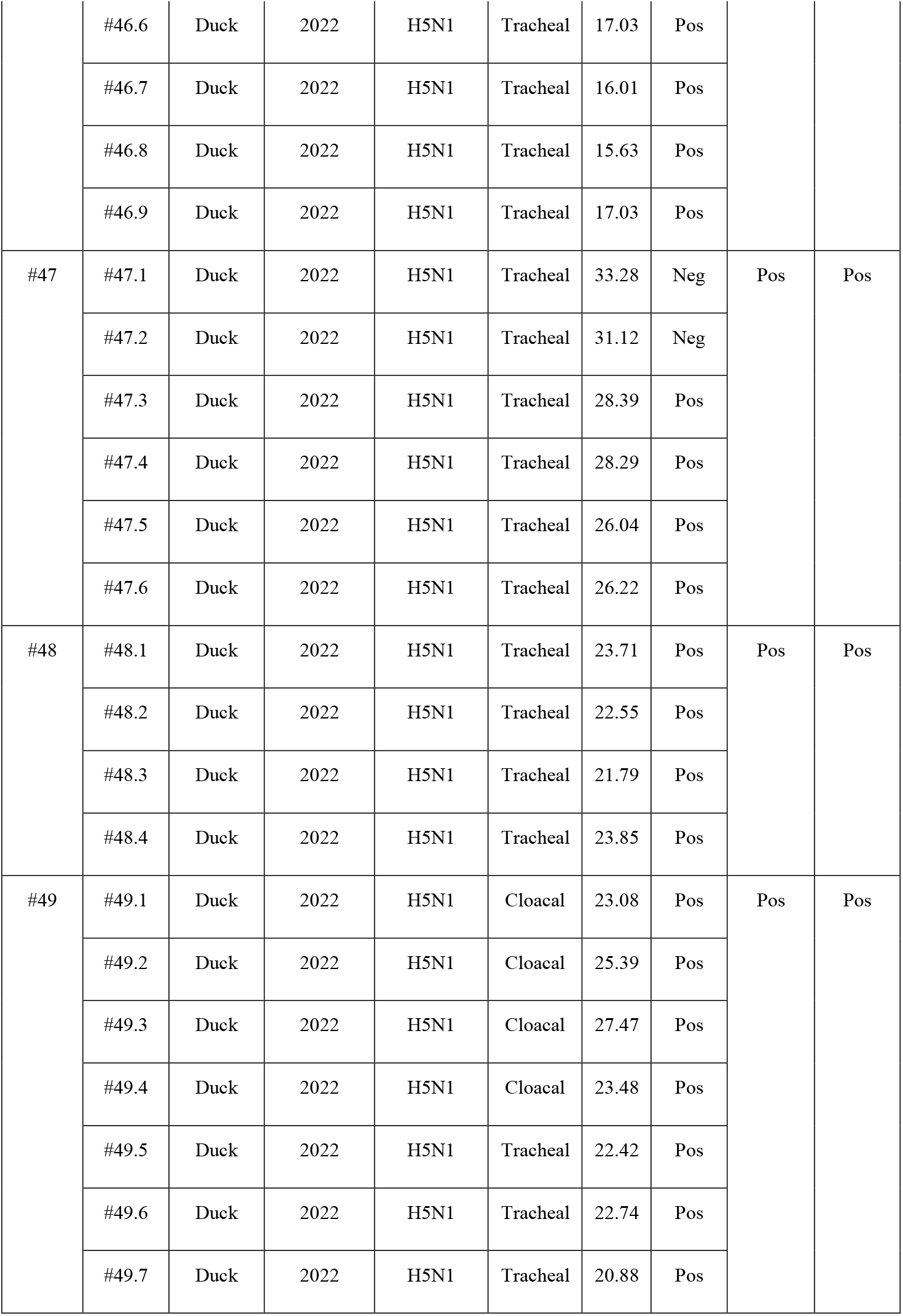
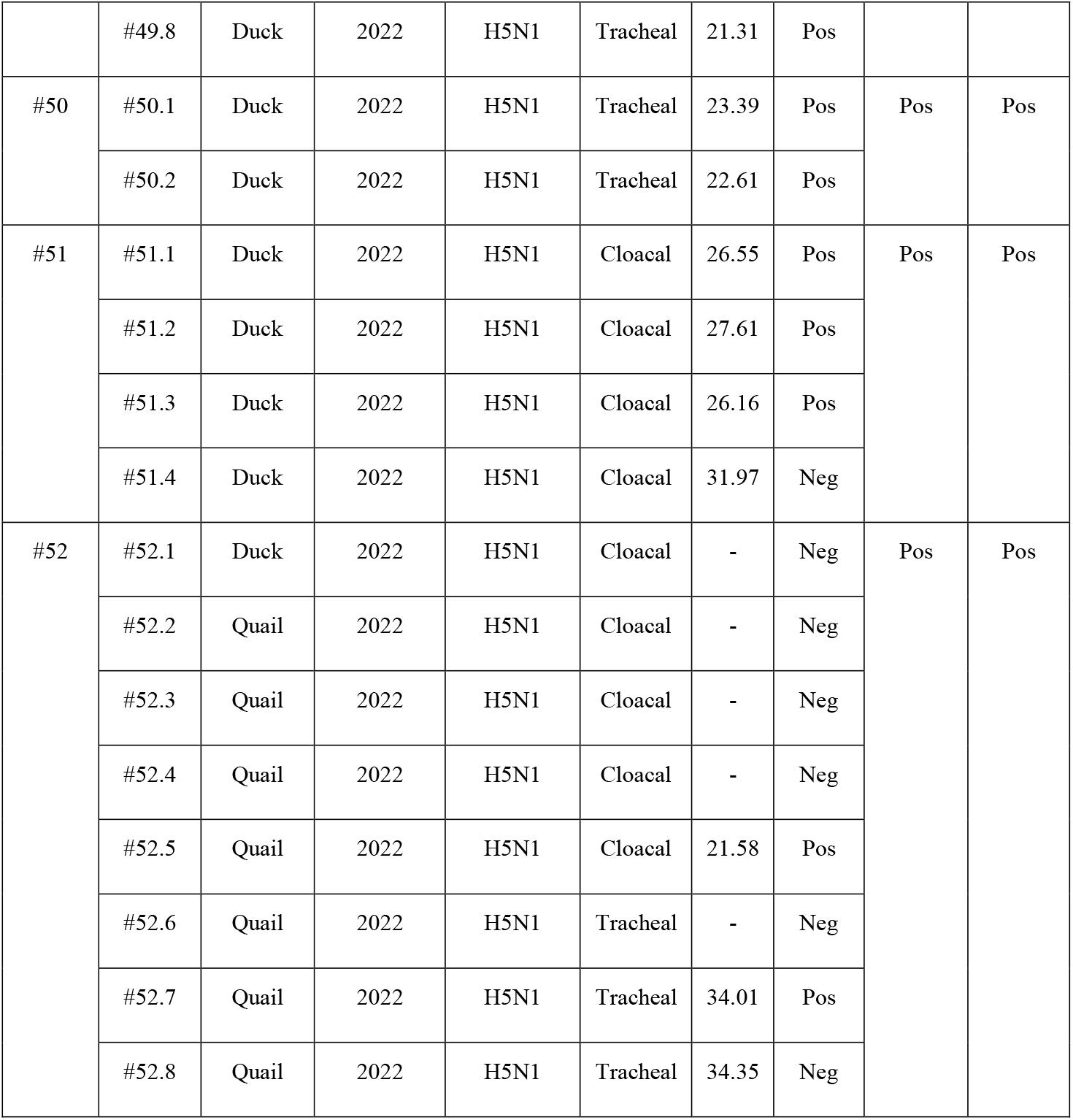
Summary of clinical sample information

**Supplementary Table 2:**
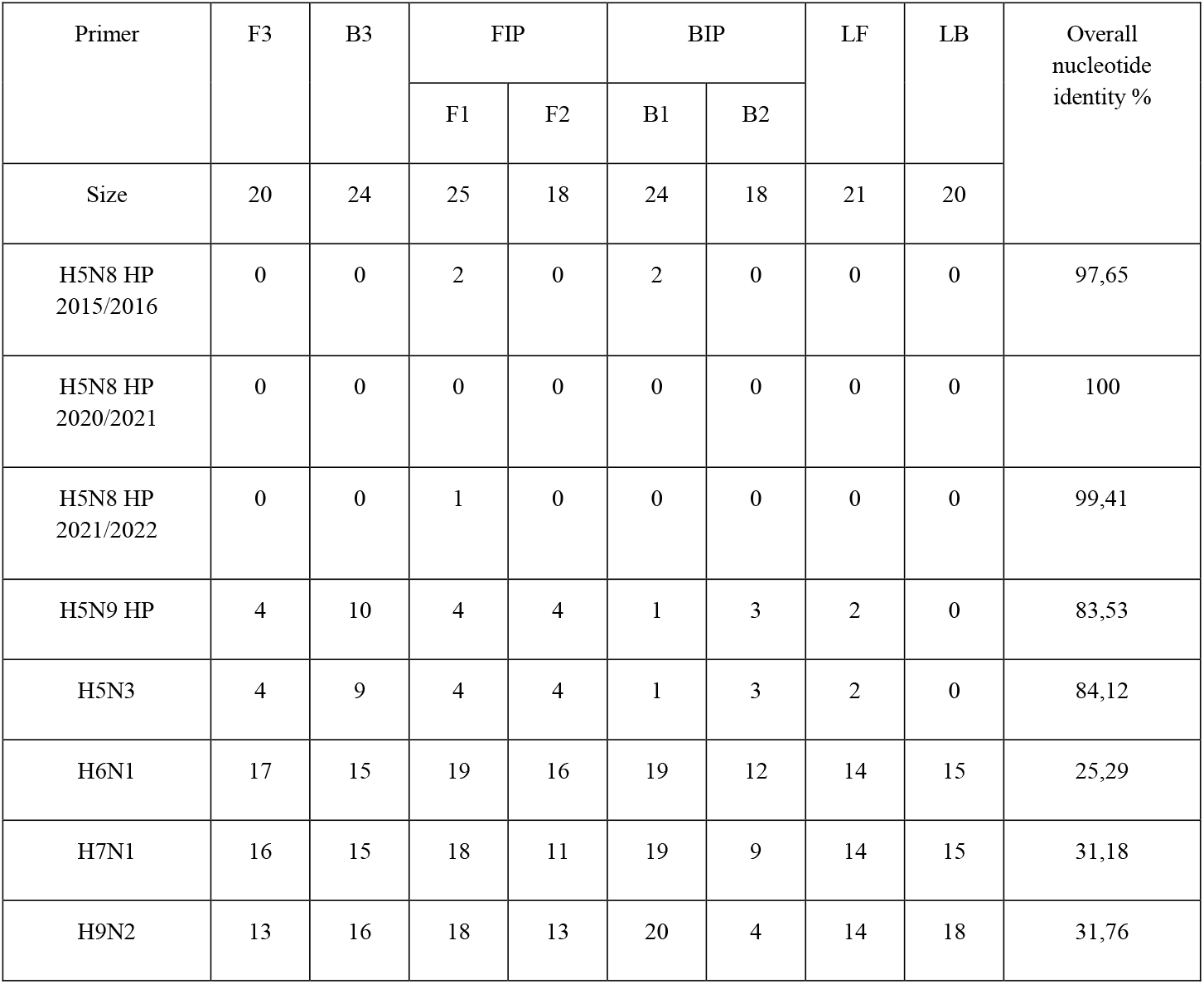
Summary of the mismatches base number between the LAMP primers and viral RNA.

| Primer | F3 | B3 | FIP |  | BIP |  | LF | LB | Overall nucleotide identity % |
| --- | --- | --- | --- | --- | --- | --- | --- | --- | --- |
|  |  |  | F1 | F2 | B1 | B2 |  |  |  |
| Size | 20 | 24 | 25 | 18 | 24 | 18 | 21 | 20 |  |
| H5N8 HP 2015/2016 | 0 | 0 | 2 | 0 | 2 | 0 | 0 | 0 | 97,65 |
| H5N8 HP 2020/2021 | 0 | 0 | 0 | 0 | 0 | 0 | 0 | 0 | 100 |
| H5N8 HP 2021/2022 | 0 | 0 | 1 | 0 | 0 | 0 | 0 | 0 | 99,41 |
| H5N9 HP | 4 | 10 | 4 | 4 | 1 | 3 | 2 | 0 | 83,53 |
| H5N3 | 4 | 9 | 4 | 4 | 1 | 3 | 2 | 0 | 84,12 |
| H6N1 | 17 | 15 | 19 | 16 | 19 | 12 | 14 | 15 | 25,29 |
| H7N1 | 16 | 15 | 18 | 11 | 19 | 9 | 14 | 15 | 31,18 |
| H9N2 | 13 | 16 | 18 | 13 | 20 | 4 | 14 | 18 | 31,76 |

**Supplementary Table 3:** HPAIV H5N8 2020/2021 and H5N1 2021/2022 RT-LAMP detection sensitivity assay.

| Dilution number |  | 1 | 2 | 3 | 4 | 5 | 6 | 7 | 8 | 9 | 10 | 11 |
| --- | --- | --- | --- | --- | --- | --- | --- | --- | --- | --- | --- | --- |
| H5N8<br>2020/2021 | rRT-PCR C<sub>t</sub> | 20.8<br>1 | 22.44 | 24.53 | 26.5<br>8 | 28.38 | 30.7<br>6 | 31.5<br>4 | 33.2<br>1 | 34.4<br>6 | 36.0<br>1 | - |
|  | RT-qPCR<br>quantification<br>copy/μL | 3327<br>0 | 1176<br>0 | 3061 | 874.<br>4 | 214.3 | 86.5<br>6 | 18.4<br>4 | 0.94 | - | - | - |
|  | Colorimetric<br>detection | Pos | Pos | Pos | Pos | Pos | Pos | Pos | Neg | Neg | Neg | Ne<br>g |
| H5N1<br>2021/2022 | rRT-PCR C<sub>t</sub> | 21.6<br>4 | 23.01 | 25.40 | 27.3<br>6 | 29.27 | 31.6<br>2 | 33 | 35.3<br>0 | 36.9<br>1 |  |  |
|  | RT-qPCR<br>quantification<br>copy/μL | 2532<br>0 | 7477 | 2424 | 665.<br>30 | 230.6<br>0 | 84.8<br>5 | 14.7<br>0 | 9.66 | - | - | - |
|  | Colorimetric<br>detection | Pos | Pos | Pos | Pos | Pos | Pos | Neg | Pos | Neg | Neg | Ne<br>g |

**Supplementary Table 4:** Overview of the LAMP results in comparison to the rRT-PCR considered as the gold standard.

| Virus subtypes | Ct range | Number of samples | True positive | True negative | False positive | False negative | Sensitivity | Specificity |
| --- | --- | --- | --- | --- | --- | --- | --- | --- |
| H5N8 HP<br>2020/2021 | Ct<25 | 59 | 59 | - | - | - | 100% | - |
|  | 25<Ct<30 | 13 | 12 | - | - | 1 | 92.31% | - |
|  | Ct>30 | 9 | 1 | 2 | - | 6 | 14.29% | 100% |
|  | All values | 81 | 72 | 2 | - | 7 | 91.14% | 100% |
| H5N1 HP<br>2021/2022 | Ct<25 | 48 | 48 | - | - | - | 100% | - |
|  | 25<Ct<30 | 38 | 32 | - | - | 6 | 84.21% | - |
|  | Ct>30 | 31 | 3 | 16 | - | 12 | 20% | 100% |
|  | All values | 117 | 83 | 16 | - | 18 | 82.18% | 100% |
| H5N8<br>&<br>H5N1 HP | Ct<25 | 107 | 107 | - | - | - | 100% | - |
|  | 25<Ct<30 | 51 | 44 | - | - | 7 | 86.27% | - |
|  | Ct>30 | 40 | 4 | 18 | - | 18 | 18.18% | 100% |
|  | All values | 198 | 155 | 18 | - | 25 | 86.11% | 100% |

